# Mechanical stress through growth on stiffer substrates impacts animal health and longevity in *C. elegans*

**DOI:** 10.1101/2024.04.11.589121

**Authors:** Maria Oorloff, Adam Hruby, Maxim Averbukh, Athena Alcala, Naibedya Dutta, Toni Castro Torres, Darius Moaddeli, Matthew Vega, Juri Kim, Andrew Bong, Aeowynn J. Coakley, Daniel Hicks, Jing Wang, Tiffany Wang, Sally Hoang, Kevin M. Tharp, Gilberto Garcia, Ryo Higuchi-Sanabria

## Abstract

Mechanical stress is a measure of internal resistance exhibited by a body or material when external forces, such as compression, tension, bending, etc. are applied. The study of mechanical stress on health and aging is a continuously growing field, as major changes to the extracellular matrix and cell-to-cell adhesions can result in dramatic changes to tissue stiffness during aging and diseased conditions. For example, during normal aging, many tissues including the ovaries, skin, blood vessels, and heart exhibit increased stiffness, which can result in a significant reduction in function of that organ. As such, numerous model systems have recently emerged to study the impact of mechanical and physical stress on cell and tissue health, including cell-culture conditions with matrigels and other surfaces that alter substrate stiffness and ex vivo tissue models that can apply stress directly to organs like muscle or tendons. Here, we sought to develop a novel method in an in vivo, model organism setting to study the impact of mechanical stress on aging, by increasing substrate stiffness in solid agar medium of *C. elegans*. To our surprise, we found shockingly limited impact of growth of *C. elegans* on stiffer substrates, including limited effects on cellular health, gene expression, organismal health, stress resilience, and longevity. Overall, our studies reveal that altering substrate stiffness of growth medium for *C. elegans* have only mild impact on animal health and longevity; however, these impacts were not nominal and open up important considerations for *C. elegans* biologists in standardizing agar medium choice for experimental assays.

## Introduction

Mechanical stress, defined as internal forces placed within a material or structure, can be classified into various types: tensile stress (stretching or pulling apart), compressive stress (squeezing or pushing together), shear stress (parallel forces acting in opposite directions), and torsional stress (twisting) [1]. In the context of living organisms, mechanical stress can have a dramatic impact on cell physiology, especially in tissues and organs subject to mechanical forces such as the muscle, bone, and blood vessels. For example, tendon and muscle tissue contain highly mechanosensitive tissue that can remodel the extracellular matrix (ECM) in response to local loading environments, which is an important response to ensure proper development, maintenance, and repair of the tissue [2]. Even on the cellular level, different substrate stiffnesses can impact cell morphology, growth, proliferation, differentiation, and matrix remodeling [3].

In response to physical forces, cells and tissues convert these mechanical forces into biochemical activities. Termed mechanochemical transduction or mechanotransduction, these events generally include extracellular receptors that initiate a signaling cascade in the cell that can regulate gene expression. For example, a wide variety of integrin receptors have been discovered that can sense changes in the extracellular environment and relay these messages inside of the cell. Changes to the extracellular matrix, physical stimulation, and alterations in cell adhesions can all activate integrin subtypes that can result in dramatic remodeling inside of the cell [4]. This includes changes to ion balance, actin and mitochondrial remodeling, and phosphorylation events that can lead to activation of genes involved in stress response, organelle homeostasis, and ECM remodeling, just to name a few [5].

Mechanical stress is also restricted to external stimuli; internal changes to tissue stiffness, such as ECM remodeling, can also initiate mechanotransduction events. This is most apparent in aging tissue, which tend to get stiffer with age due to a build-up of collagen that can contribute to tissue fibrosis [6]. This increase in tissue stiffness can have direct consequences on tissue function. For example, changes in collagen composition and cross-linking in the tendon with age can result in reduced tendon compliance, increased stiffness, and decreased resilience to mechanical stress, which is the direct cause of age-related increase in tendon injuries [7,8]. In addition, the aging ovary displays a collagen-dependent increase in stiffness, which results in improper follicle development and oocyte quality and an overall reduction in reproductive quality. These changes were not only limited to collagen, but also involve changes to other components of the ECM, including a significant decrease in the glycosaminoglycan, hyaluronic acid (HA) [9]. In fact, HA is a commonly used anti-aging intervention for skin health as it can regulate skin moisture and elasticity [10], and even in cell culture models and model organisms, increased levels of HA can improve stress resilience and longevity [11].

Changes to ECM composition not only occur during aging but are a hallmark of cancer pathology. Solid state tumors are characterized by a significant increase in stiffness due to increased density of collagen and other ECM components [12]. This change in tumor microenvironment can drive cancer progression, as increased adhesion-mediated mechanosignaling can initiate “pro-survival” mechanisms in cancer cells. Specifically, one study found that cancer cells growing on stiffer substrates activate an integrin-mediated remodeling of the cytoskeleton and mitochondria, which can activate cellular stress responses that improve cancer cell resilience [13]. This increase in stress resilience translated to increased drug-resistance, which can be ameliorated by downregulating these pathways, suggesting that targeting mechanotransduction pathways can have potential both in anti-aging and anti-cancer interventions.

Considering the significance of mechanotransduction in organismal health and aging, we sought to determine the effect of mechanical stress on an in vivo model system, *C. elegans*. Ideally, our goal was to identify the simplest method to apply mechanical stress onto this model organism in an effort to develop a model system that could be used to study how changes in mechanotransduction can impact the aging process. As such, we applied mechanical stress by growing *C. elegans* on a stiffer substrate, which previous studies have shown can alter *C. elegans* locomotory behavior [14]. Here, we created a stiffer substrate by using double the concentration of agar (4%) compared to a standard solid agar medium (2%). Although we expected this doubling of substrate stiffness to have major impacts on organismal health, our work revealed that growth of animals on stiffer substrates did not activate canonical mechanotransduction pathways. Moreover, cellular health and transcriptional regulation were wildly unaffected, which resulted in shockingly mild – if any – changes to whole organism health, stress resilience, and longevity.

## Results

To develop a mechanical stress model for *C. elegans*, here we sought to alter substrate stiffness by growing animals on higher percentages of agar. Our standard lab-growth nematode growth medium (NGM) contains 2% agar w/v and for the entirety of this study, we used Bacto Agar Solidifying Agent from BD Diagnostics (catalog no. 281210). For our stiffer substrate, we utilized 4% agar w/v, as this was the highest concentration of agar we could accomplish without having issues with the agar either precipitating or solidifying prior to cooling sufficiently to add liquid additives for standard NGM plate-pouring (refer to [15] for a detailed protocol on making NGM plates). When wild-type animals were grown on stiffer substrates from the L1 larval stage, we found that animals exhibited a mild, but significant increase in lifespan (**Fig. 1A**).

**Fig. 1.**
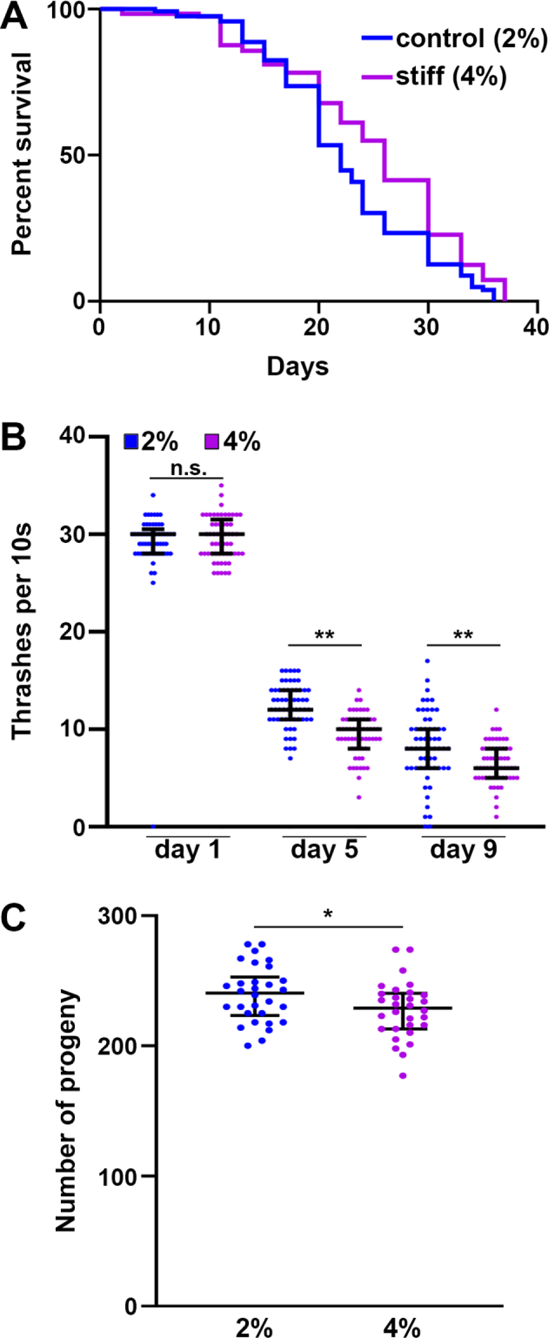
Growth on stiff substrate results in mild lifespan extension but decrease in organismal health. **(A)** N2 wild-type animals grown on empty vector (EV) RNAi bacteria on either control (2%, blue) or stiff (4%, purple) agar plates from L1. Lifespans were scored every 2 days. Data is representative of 3 replicates and statistical analysis is available in **Table S1**. **(B)** N2 wild-type animals were grown on EV RNAi bacteria on either 2% or 4% NGM plates from L1. At day 1, 5, and 9 of adulthood, animals were collected in M9 media and video recordings were taken on an M205 stereoscope for 10s. Body bends per 10s were counted by eye for each individual worm. **(C)** N2 wild-type animals were grown on EV RNAi bacteria on either 2% or 4% agar plates from L1. At the L4 stage, animals were singled out and moved daily. Number of live progenies were counted for each animal with n > 50 per sample. Data is pooled from 3 independent biological replicates with n = 10 per replicate. * = p < 0.05, ** = p < 0.01 through non-parametric Mann-Whitney testing. Lines are median and interquartile range, and each dot represents a single animal where blue dots are animals grown on 2% control plates and purple dots are animals grown on 4% stiff plates.

Interestingly, although animals live slightly longer, they display a mild decrease in specific healthspan metrics, including locomotory behavior measured by a liquid thrashing assay (**Fig. 1B**) and reproductive capacity (**Fig. 1C**). Therefore, although animals are slightly longer-lived when grown on stiffer substrates, there appears to be a tradeoff in healthspan.

Here, we reasoned that growth on stiffer substrates may extend lifespan by activation of specific stress response pathways. Indeed, numerous previous studies have shown hyperactivation of specific stress responses, including the unfolded protein responses of the mitochondria (UPR^MT^) [16], the endoplasmic reticulum (UPR^ER^) [17], and heat-shock response (HSR) [18] all extend longevity. In addition, several of these stress response-mediated longevity paradigms are associated with a decrease in reproductive potential [15,19], which mirrors phenotypes found on animals grown on stiffer substrates. However, we found that stress responses were largely unchanged (**Fig. 2**). Specifically, we measured activation of stress responses using transcriptional reporters whereby GFP expression is under the promoter of canonical genes activated upon stress. For UPR^MT^, the promoter of the mitochondrial chaperone, *hsp-6* (HSPA9 in mammals) [20] is used. Growth on 4% agar had no impact on *hsp-6p::GFP* expression, both in the presence (RNAi knockdown of the electron transport chain component, *cco-1* [16]) and absence of stress (**Fig. 2A**). Similarly, we measured UPR^ER^ induction using the *hsp-4p::GFP* (HSPA5/BiP) reporter [21] and found no activation of UPR^ER^ upon growth on stiffer substrates. There was a very mild increase in UPR^ER^ expression under stress applied by RNAi knockdown of *tag-335* involved in N-linked glycosylation of proteins in the ER, knockdown of which drives accumulation of misfolded proteins in the ER. However, similar to UPR^MT^, there were no measurable changes in HSR using the *hsp-16.2p::GFP* (CRYAB) reporter [22] or oxidative stress response using the *gst-4p::GFP* (HPGDS) reporter [23] both in the presence (2-hour heat shock at 34°C or tert-butyl hydroperoxide exposure) and absence of stress (**Fig. 2C-D**).

**Fig. 2.**
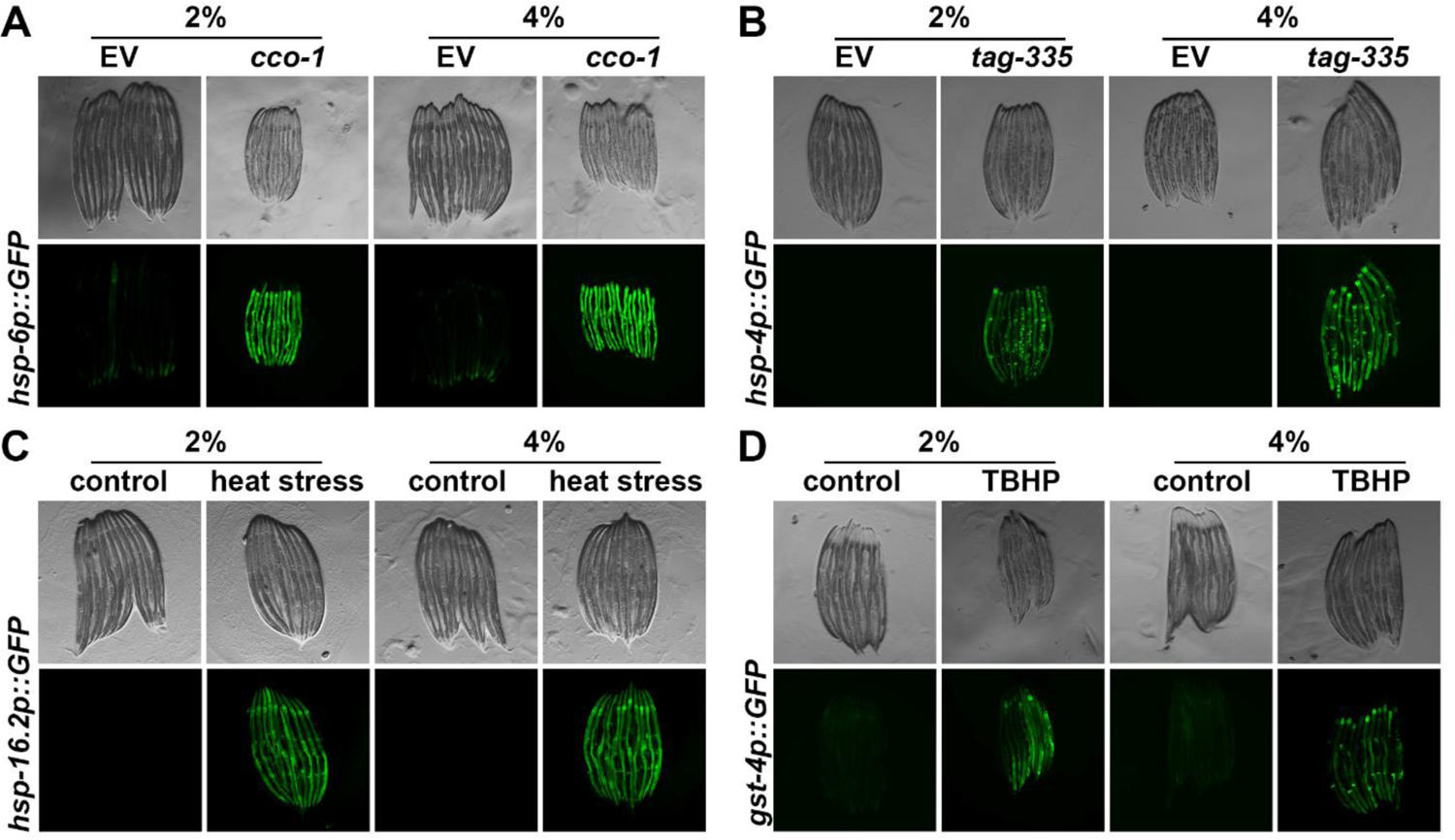
Growth on stiff substrates does not affect induction of stress responses. **(A)** Representative fluorescent images of day 1 adult animals expressing *hsp-6p::GFP* grown on EV or *cco-1* RNAi bacteria from L1. Data is representative of 3 independent replicates. **(B)** Representative fluorescent images of day 1 adult animals expressing *hsp-4p::GFP* grown on EV or *tag-335* RNAi bacteria from L1. Data is representative of 3 independent replicates. **(C)** Representative fluorescent images of day 1 adult animals expressing *hsp-16.2p::GFP* grown on EV RNAi bacteria from L1. Animals were heat-shocked at 34°C for 2 hours followed by a 2 hour recovery at 20°C. Data is representative of 3 independent replicates. **(D)** Representative fluorescent images of day 1 adult animals expressing *gst-4p::GFP* grown on EV RNAi bacteria from L1. Animals were treated with 1 mM tert-butyl hydroperoxide (TBHP) rotating in an M9-TBHP solution for 2 hours at 20°C. TBHP was then washed with M9 and worms were recovered on OP50 bacteria for 16 hours prior to imaging. Data is representative of 3 independent replicates.

Although transcriptional reporters provide a robust and rapid method of screening for activation of stress responses, a major limitation is that they focus on a single target, whereas stress responses tend to involve major and dramatic remodeling of transcription. Therefore, to obtain a more comprehensive overview of transcriptional changes driven upon growth on stiffer substrates, we performed bulk RNA-seq analysis comparing animals grown on standard 2% and stiffer 4% agar plates (**Fig. 3**). Shockingly, we saw very minor changes in transcription, with only 35 genes differentially expressed (with an adjusted p < 0.05 cutoff) in animals grown on stiffer substrates (**Fig. 3A-B**). Importantly, very few of these gene expression changes were those found in canonical mechanical stress or mechanotransduction pathways, including remodeling of the ECM, mitochondria, or actin cytoskeleton. Only two GO-terms associated with extracellular areas of the cell were identified (**Fig. 3C**). Instead, the primary change we saw were those associated with the vacuole and lysosome, including *lipl-1*, encoding a key lysosomal lipase involved in lipophagy [24] and *asm-3*, encoding a sphingomyelinase involved in sphingomyelin breakdown, potentially through the lysosome [25]. To determine whether these changes in expression of lysosome and lipid-recycling related genes had direct physiological impact, we next measured lysosome function and lipid droplet content. Interestingly, we did not observe any major differences in lysosomal quantity using LMP-1::GFP [26] (**Fig. S1A**), nor did we see any changes in lipid droplets as measured using DHS-3::GFP [27] (**Fig. S1B**).

**Fig. 3.**
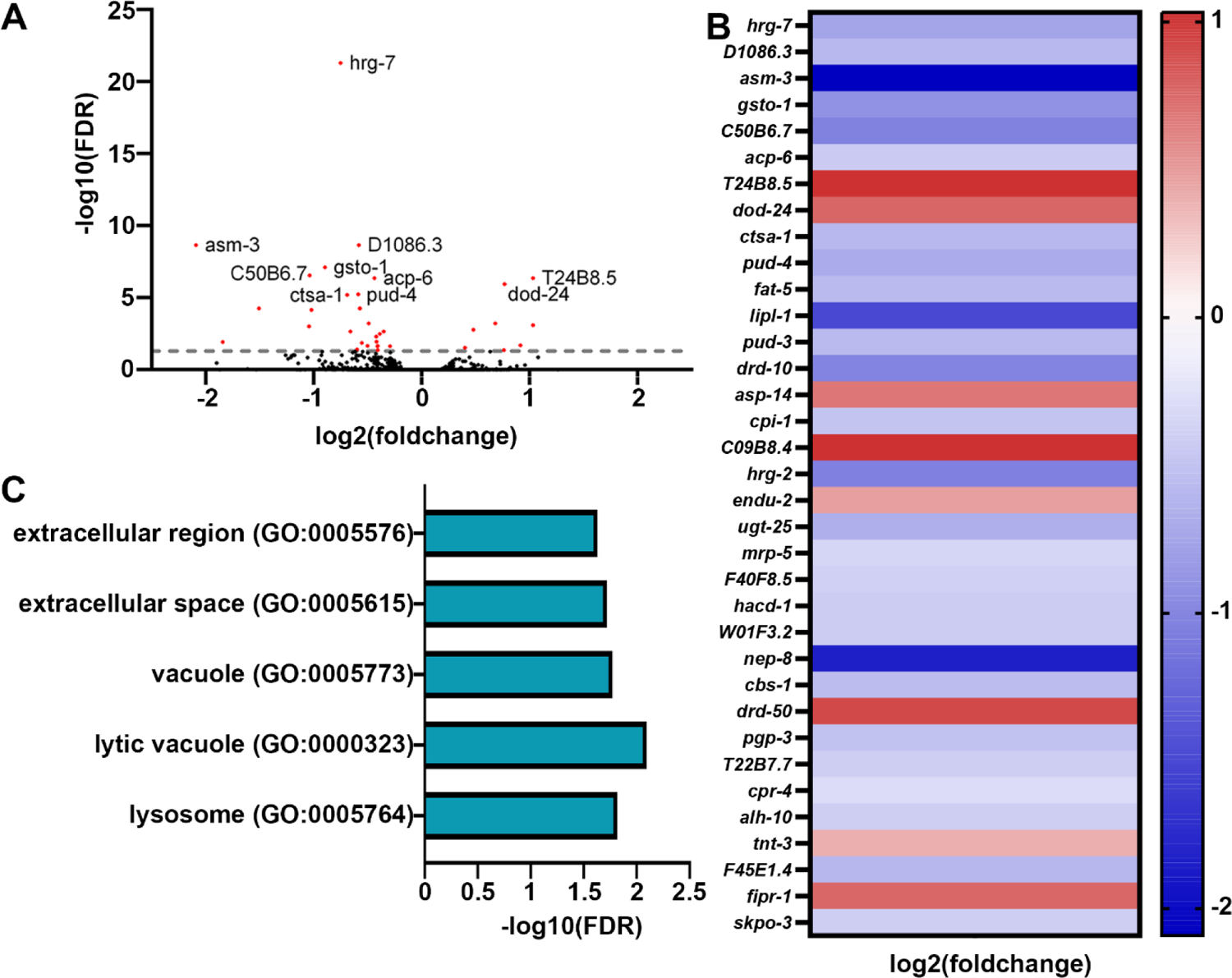
Transcriptome analysis reveals minor changes in gene expression in animals grown on stiffer substrates. **(A)** Volcano plot of differentially expressed genes of animals grown on stiff (4%) agar compared to standard (2%) agar. Every dot is a single gene, where red dots indicate genes with p-value < 0.05 and black dots indicate genes with p-value > 0.05. **(B)** Heat map of all differentially expressed genes in worms grown on 4% agar where warmer colors indicate higher expression and cooler colors indicate lower expression. A list of all genes are available in **Table S2**. **(C)** Gene ontology enrichments for differentially expressed genes (*p-value* < 0.05) in worms grown on 4%’ agar.

Since we saw a slight increase in UPR^ER^ activation under conditions of stress in animals grown on stiffer substrates, we next sought to determine whether these animals have an increase in a functional and beneficial UPR^ER^. To test this, we grew animals on tunicamycin, which induces ER stress and significantly decreases lifespan [17]. Interestingly, we find that animals grown on stiffer substrates have a significant increase in survival under exposure to tunicamycin, which was dependent on the main UPR^ER^ transcription factor XBP-1, as knockdown of *xbp-1* suppressed this increase in resilience (**Fig. S2A**). Together, these data suggest that animals grown on stiffer substrates likely exhibit a functional increase in UPR^ER^ activity upon stress exposure, which is dependent on canonical induction of UPR^ER^ through XBP-1 [28]. Importantly, this is specific to ER stress resilience as animals on stiffer substrates did not exhibit an increase in oxidative or mitochondrial stress as measured by survival upon exposure to paraquat (**Fig. S2B**), which induces superoxide formation in the mitochondria [29].

Previous studies have shown that exposure to ER stress can result in dramatic remodeling of the ER and activation of lipophagy machinery [30]. Since we saw an increase in ER stress resilience, we next visualized ER morphology using an mRuby::HDEL fusion protein that localizes to the ER lumen through a SEL-1 signal sequence [30]. However, we did not see any major differences in ER morphology (**Fig. S3A**).

To get a better sense of wholescale cellular changes upon growth on stiffer substrates, we expanded our studies to changes in mitochondria and actin organization, as previous research has shown that growth on stiffer substrates can result in integrin-dependent remodeling of the mitochondria and actin cytoskeleton in both human cells [13] and *C. elegans* [31]. Here, we visualized mitochondrial morphology in the muscle, intestine, and hypodermis of *C. elegans* using a mitochondria-targeted GFP [32] and cell-type specific effects on mitochondrial morphology (**Fig. S3B-D**). We find that mitochondria were fragmented in muscle, intestine, and the hypodermis in animals grown on stiffer substrates. To measure actin organization, we used the F-actin binding protein, LifeAct::mRuby expressed specifically in the muscle, intestine, and hypodermis [33]. As previously reported, the actin cytoskeleton shows marked deterioration visualized by decreased linear organization of actin filaments in the muscle [33]. Interestingly, we saw an increase in actin stability at advanced age in animals grown on stiffer substrates (**Fig. 4A**). These data are consistent with previous studies that showed that increased integrin signaling can prevent age-related decline in actin organization, which can also drive mitochondrial fragmentation [31].

**Fig. 4.**
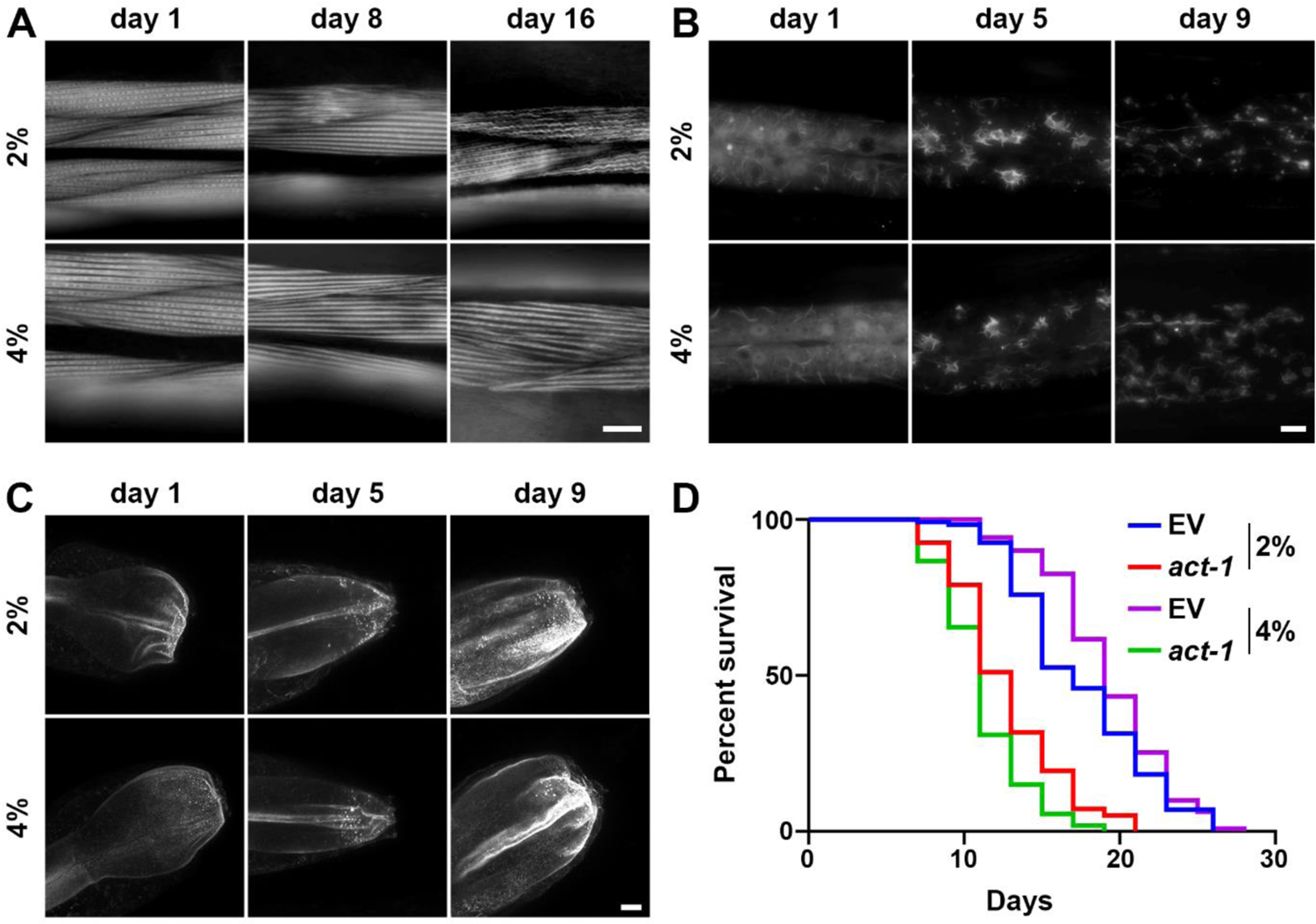
Growth on stiff surfaces results in increased actin stability, which can drive lifespan extension. **(A)** Representative fluorescent images of body wall muscle actin (*myo-3p::LifeAct::mRuby*) are shown. Animals were grown on empty vector (EV) RNAi from L1. Single-slice images were captured on a Leica THUNDER Imager. Scale bar is 10 µm. **(B)** Representative max-projection fluorescent images of hypodermal actin (*col-19p::LifeAct::mRuby*) are shown. Animals were grown on empty vector (EV) RNAi from L1. Z-stack images were captured on a Leica THUNDER Imager using system-optimized z-slices. Scale bar is 10 µm. **(C)** Representative max-projection fluorescent images of intestinal actin (*gly-19p::LifeAct::mRuby*) are shown. Animals were grown on empty vector (EV) RNAi from L1. Z-stack images were captured on a Leica Stellaris using system-optimized z-slices. Scale bar is 10 µm. **(D)** N2 wild-type animals grown on empty vector (EV) or *act-1* RNAi diluted 10/90 *act-1*/EV on either control (2%) or stiff (4%) agar plates from L1. Lifespans were scored every 2 days. Data is representative of 3 replicates and statistical analysis is available in **Table S1**.

The hypodermis displays rearrangement of the actin cytoskeleton to star-like structures that are likely endocytic vesicles at around day 3-5 of adulthood, which start to dissociate after day 7 [33]. Similar to muscle actin, growth on stiffer substrates results in a mild increase in stability of actin, such that more structures are preserved at late age (**Fig. 4B**). Finally, actin in the intestine exhibits mislocalization and aggregation, which can result in decreased gut barrier function during aging [34]. Interestingly, we do not see improvements of these phenotypes in the intestine of animals grown on stiffer substrates (**Fig. 4C**). To determine whether the mild increase in these animals is associated with the corresponding increase in actin stability, we disrupted actin function using a carefully titrated, non-lethal dosage of actin RNAi as previously described [35]. Remarkably, we found that low-dose actin RNAi complete suppressed the lifespan extension found in these animals (**Fig. 4D**). In addition, animals grown on stiff substrates displayed a slight increase in maximum thermotolerance (**Fig. S4**). Although in general, median thermotolerance did not show a significant increase, each replicate did display a significant difference from the 2% control (see **Table S1** for statistical analysis). Importantly, an increase in thermotolerance has previously been ascribed to an increase in cytoskeletal stability [36]. These data suggest that growth on stiffer substrates does result in an increase in actin stability, which is likely responsible for the mild increase in longevity found in these animals.

## Discussion

Mechanotransduction represents the complex interplay between the extracellular environment and the internal biological system. All living organisms are exposed to environmental changes that can apply mechanical forces, which are then converted into biochemical activities through a signaling cascade involving extracellular receptors and intracellular signaling pathways, culminating in remodeling of numerous intracellular components on the gene, protein, and organelle levels. For example, in human breast cancer cell lines, growth on stiffer substrates results in alterations of mitochondrial composition, structure, and function. These changes are driven by integrin-mediated mechanotransduction signals that alter solute transporters, including SLC9A1, which results in transcriptional activation of HSF1 to alter actin and mitochondrial function and oxidative stress response [13]. These dramatic intracellular changes both on the transcriptional and organellar level occur in response solely to alterations in substrate stiffness and have direct ramifications for cancer cell survival.

Here, we tested the impact of growing whole animals on stiffer substrates. Specifically, the nematode *C. elegans* was grown on NGM containing 4% agar compared to the standard 2% agar concentration, which results in a stiffer plate and recapitulates a handful of phenotypes found in in vitro studies. First, animals exhibited an increase in actin stability, which is consistent with previous studies that showed that increased integrin signaling in *C. elegans* results in increased stability of actin filaments at late age [31]. Moreover, this improvement in actin stability plays an important role in increasing longevity, again consistent with several studies that have linked actin function and stability to lifespan in *C. elegans* [33,35]. Despite this actin-dependent increase in lifespan, animals grown on stiffer substrates did not exhibit major changes in gene expression of actin-related genes. It is possible that regulation of actin downstream of exposure to stiffer substrates is not dependent on transcriptional changes, but rather to protein signaling events, such as those mediated by Rho GTPases, an important actin-regulatory signaling switch downstream of mechanosensing [37]. Indeed, actin remodeling downstream of increased integrin signaling in human cells is dependent on RhoA/RAC signaling [13]. However, the lack of a general mechanotransduction gene expression signature suggests that perhaps whole animals grown on stiffer substrates result in activation of different signaling pathways than cells grown directly on stiffer substrates. For example, it is possible that the thick cuticle of *C. elegans* – the part of the whole animal that interacts with the agar substrate – has dramatic differences in mechanosensing and downstream mechanotransduction compared to cells.

In addition to a lack of changes in actin-related genes, our RNA-seq analysis also did not show changes in UPR^MT^ or other mitochondrial stress-related pathways. This was surprising considering the fragmentation of mitochondria observed in animals grown on stiff substrates and the activation of UPRMT pathways visible in other stiffness and integrin-related paradigms [13,31]. Perhaps animals grown on stiffer substrates may not activate integrin signaling to a sufficient level to induce transcriptional changes but do increase mechanosignaling pathways enough to promote actin quality, which can have an impact on mitochondrial morphology. This is an important distinction as animals grown on stiffer substrates did not exhibit an increase in resilience to mitochondrial or oxidative stress, whereas animals with increased integrin signaling or cells grown on stiffer substrates in vitro did exhibit a significant increase in both mitochondrial and oxidative stress resilience [13,31]. Moreover, animals with increased UPR^MT^ signaling also exhibit an increase in resistance to paraquat [38], adding further evidence that animals grown on stiffer substrates do not activate a canonical UPR^MT^ pathway.

Although changes to the transcriptome were limited, a few lysosomal genes related to lipid metabolism, including lysosomal lipases, were identified. A previous study showed that downstream of UPR^ER^ activation, increased lysosomal lipases can drive increased ER stress resilience and longevity [39]. This is an important consideration as animals grown on stiffer substrates also exhibit an increase in ER stress resilience, which is dependent on the UPR^ER^ regulator, *xbp-1s*. While these data may suggest that the increase in lysosomal lipases and the associated increase in ER stress resilience may drive the lifespan extension found in these animals, we did not exhibit global changes to lysosome levels or the autophagy marker LGG-1. In addition, we also did not see any changes to whole animal lipid levels. Thus, further studies are required to determine whether these mild changes in lysosome-related genes identified in our study are sufficient to drive ER stress resilience and longevity phenotypes.

Even more interesting would be to determine whether there are any overlaps between the ER and actin-related phenotypes. Indeed, a recent study has shown that promoting actin stability by increased expression of the chromatin remodeling factor, BET-1, has direct impact on ER stress resilience [35]. Moreover, the mammalian homologue of BET-1, BRD4, has been linked to pathological fibrosis in multiple tissues, including the cornea [40] and lung [41], likely through regulation of mechanosignaling. Thus, actin and ER regulation may have significant overlap, which can potentially be downstream of mechanotransduction pathways.

Overall, our study illustrated that whole animals grown on stiffer substrates do have general changes to cell biology, physiology, healthspan, and lifespan, although these effects are quite moderate. Importantly, several of these changes are correlated with changes seen in other models of mechanotransduction, although not all pathways and changes seen in other systems were observed in our animals. Although the phenotypes presented in this study may be mild and may not fully recapitulate what is expected for a mechanotransduction model, there is an important lesson here for *C. elegans* biologists. It is imperative that NGM agar plates not be stored long-term, as desiccation of plates can indirectly increase agar precentage relative to water content. These plates will be stiffer and can change your biological data, especially for phenotypes similar to those we assayed in this study. Another very important consideration is to standardize an agar choice in the lab since multiple different agar sources have dramatically different stiffnesses (**Fig. 5**). While our study exemplifies the importance of standardizing an agar choice for *C. elegans* biology, it is possible that this consideration is important for any study involving agar-based solid medium.

**Fig. 5.**
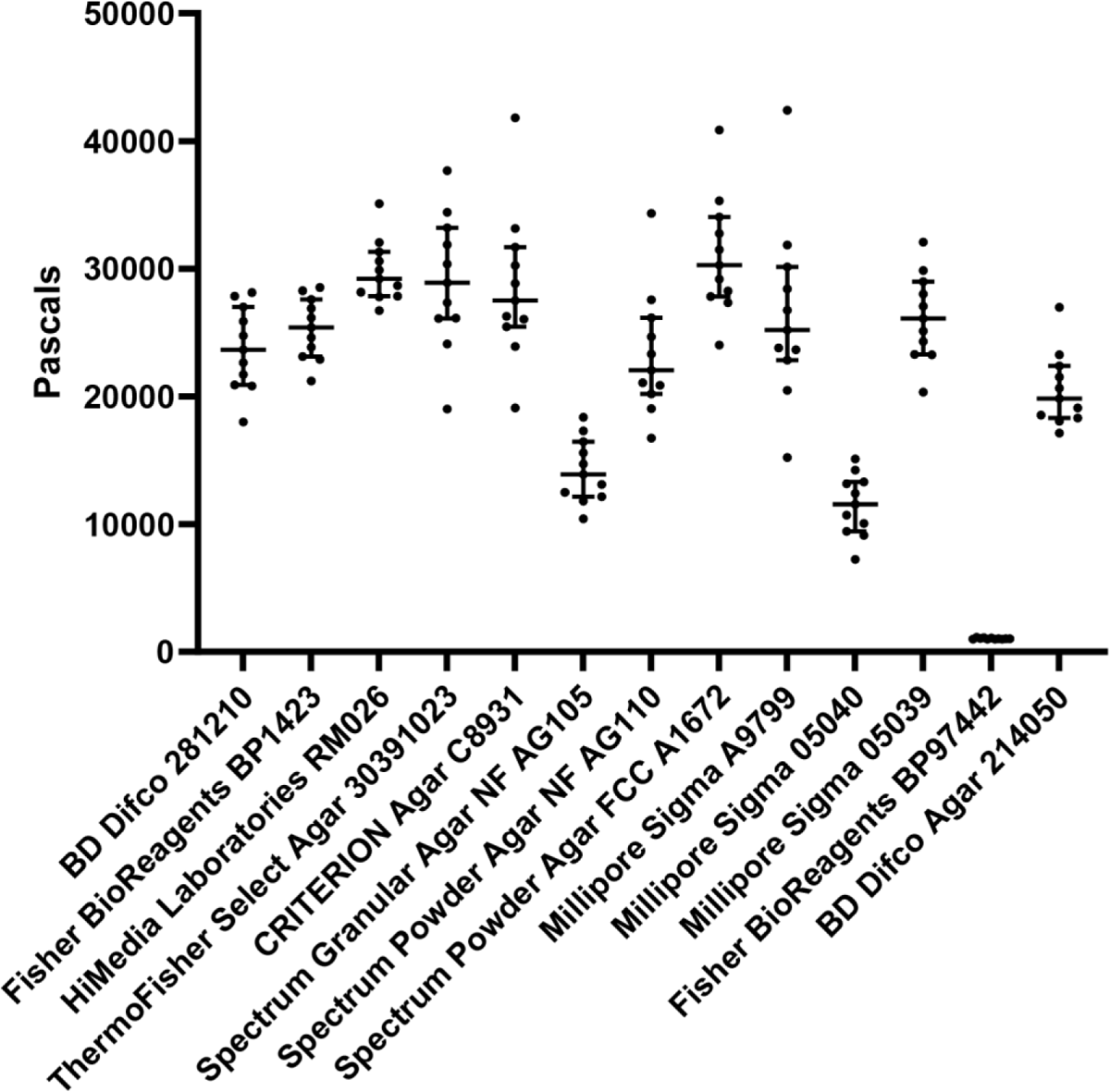
Variable sources of agar formulations have significant differences in stiffness. Standard 2% agar-based solid NGM plates were made as described in Materials and Methods using different sources of agar. Stiffness of each agar plate was measured using an oscillatory rheometer. Each dot represents a single technical replicate performed on an individual plate and lines represent median plus interquartile range. X-axis indicates brand and catalog number of each agar source.

## Materials and Methods

### *C. elegans* maintenance

All strains used in this study are derivatives of the N2 wild-type strain from the Caenorhabditis Genetics Center (CGC). All animals were grown at 15°C on OP50 *E. coli* B strain and moved onto a new NGM plate seeded with OP50 every week for standard maintenance. To avoid genetic drift, every week > 20 individual animals were moved to new plates each passage for a maximum of 25-30 passages. All experimental strains were bleached to clarify OP50 bacteria and grown at 20°C on HT115 E. *coli* K strain bacteria. For all experiments, animals were either grown on HT115 bacteria carrying a pL4440 plasmid that does not target any specific genes (called EV for empty vector in this manuscript) or pL4440 vector carrying a partial gene sequence against a specific target gene for RNAi-mediated knockdown.

### Plates

Standard NGM plates for maintenance: Bacto-Agar (Difco) 2% w/v, Bacto-Peptone 0.25% w/v, NaCl2 0.3% w/v, 1 mM CaCl2, 5 µg/ml Cholesterol, 0.625 mM KPO4 pH 6.0, 1 mM MgSO4. 2% RNAi plates for experiments: Bacto-Agar (Difco) 2% w/v, Bacto-Peptone 0.25% w/v, NaCl2 0.3% w/v, 1 mM CaCl2, 5 µg/ml Cholesterol, 0.625 mM KPO4 pH 6.0, 1 mM MgSO4, 100 µg/mL Carbenicillin, 1 mM IPTG. 4% RNAi plates for experiments: Bacto-Agar (Difco) 4% w/v, Bacto-Peptone 0.25% w/v, NaCl2 0.3% w/v, 1 mM CaCl2, 5 µg/ml Cholesterol, 0.625 mM KPO4 pH 6.0, 1 mM MgSO4, 100 µg/mL Carbenicillin, 1 mM IPTG. For all aging experiments, 100 uL of 10 mg/mL FUDR was placed directly on the bacterial lawn.

### Bleaching

All experiments were performed on synchronized, age-matched populations acquired through a standard bleaching protocol. Animals were collected into a 15mL conical tube using M9 solution (22mM KH2PO4 monobasic, 42.3mM NaHPO4, 85.6 mM NaCl, 1 mM MgSO4). M9 solution was replaced with bleaching solution (1.8% sodium hypochlorite, 0.375 M KOH, diluted in M9 solution) and animals are treated with bleaching solution until only eggs remain in the tube with no adult carcasses remaining (∼4-6 minutes). Eggs were then washed with M9 solution 3x by centrifugation at 1.1 RCF for 30 seconds and aspiration of bleaching solution and replacing with M9 solution. Following the final wash, animals were L1 arrested in M9 solution in a rotator for up to 24 hours in 20°C.

| Strains used in this study |  |  |
| --- | --- | --- |
| *C. elegans*: Bristol (N2) strain as wild type (WT) | CGC | N2 |
| *C. elegans*: CL2070: *dvls70*[*pCL25 (hsp-16.2p::GFP)*, *pRF4(rol-6)*] | CGC |  |
| *C. elegans*: CL2166: *dvls19*[*pAG15(gst-4p::GFP::NLS)*] III | CGC |  |
| *C. elegans*: AGD1664: *uthSi17*[*myo-3p::MLS::GFP::unc-54 3'UTR::cb-unc-119(+)*] I; *unc-119(ed3)* III | [32] |  |
| *C. elegans*: AGD1988: *zcls13*[*hsp-6p::GFP*] | [35] | SJ4100 6x backcross |
| *C. elegans*: AGD2053: *zcls4*[*hsp-4P::GFP*]V | [42] |  |
| *C. elegans*: AGD2192: *unc-119(ed3)* III; *uthSi60*[*vha-6p::ERss::mRuby::HDEL::unc-54 3'UTR cb-unc-119(+)*] IV; AGD2192: | [30] |  |
| *C. elegans*: DA2123: *adls2122*[*lgg-1p::GFP::lgg-1 + rol-6(su1006)*] | CGC |  |
| *C. elegans*: RT258: *pwls50*[*Imp-1::GFP + Cbr-unc-119(+)*] | CGC |  |
| *C. elegans*: RHS10: *ldrls*[*dhs-3p::dhs-3::GFP + unc-76(+)*] | This study | LIU1 6x backcross |
| *C. elegans*: RHS41: *uthSi7*[*myo-3p::LifeAct::mRuby::unc-54 3'UTR::cb-unc-119(+)*] IV | [33] | AGD1651 4x backcross |
| *C. elegans*: RHS42: *uthSi10*[*col-19p::LifeAct::mRuby::unc-54 3'UTR::cb-unc-119(+)*] IV | [33] | AGD1654 4x backcross |
| *C. elegans*: RHS43: *uthSi13*[*gly-19p::LifeAct::mRuby::unc-54 3'UTR::cb-unc-119(+)*] IV | [33] | AGD1657 4x backcross |

### Microscopy

#### Steromicroscope

For transcriptional reporter imaging (*hsp-6p::GFP, hsp-4p::GFP, hsp-16.2p::GFP, gst-4p::GFP)*, synchronized animals were grown on either 2% RNAi or 4% RNAi plates from L1. Animals were imaged at day 1 of adulthood and for aging experiments also imaged at day 5 and day 9 of adulthood. Animals were placed on standard NGM plates without bacteria on 100 mM sodium azide to paralyze worms. Paralyzed animals were lined up alongside each other and imaged on a Leica M205FCA automated fluorescent stereomicroscope equipped with a standard GFP and RFP filter, Leica LED3 light source, and Leica K5 camera and run on LAS X software. For all imaging experiments, three biological replicates were performed, and one representative replicate was chosen for the figures.

#### Widefield and confocal imaging

For high resolution imaging of organelles including actin, mitochondria, ER, lysosomes, and lipid droplets, we performed imaging on a Leica THUNDER widefield microscope equipped with a 63x/1.4 Plan AproChromat objective, standard GFP and dsRed filter, Leica DFC9000 GT camera, a Leica LED5 light source, and run on LAS X software or a Leica Stellaris 5 confocal microscope equipped with a white light laser source and spectral filters, HyD detectors, 63x/1.4 Plan ApoChromat objective, and run on LAS X software. Animals were placed in 100 mM sodium azide solution on a glass slide and imaged within 5 minutes of slide preparation to prevent artifacts from animals being on a glass slide in sodium azide for extended periods of time. For all mitochondrial morphology imaging, no sodium azide was used, and animals were placed on a glass slide with M9 solution and imaged within 5 minutes to prevent mitochondrial fragmentation. For all imaging experiments, three biological replicates were performed, and one representative replicate was chosen for the figures.

### RNAseq analysis

RNA isolation was performed on day 1 of adulthood. ∼1000 animals were grown on either 2% or 4% RNAi plates with HT115 bacteria and animals were harvested from plates using M9.

Animals were gravity settled to allow adult worms to settle on the bottom and eggs/L1s were aspirated off the top. Animals were washed 3x in this manner to minimize the number of progeny collected. Animals were then placed into Trizol solution and worms were freeze/thawed 3x with cycles of liquid nitrogen and 37 °C water bath for 1 min. Before every freeze cycle, animals were vortexed for 30 s to ensure complete digestion of animal cells. After the final thaw, chloroform was added at a 1:5 chloroform:trizol ratio and aqueous separation of RNA was performed via centrifugation in a heavy gel phase-lock tube (VWR, 10847-802). The aqueous phase was mixed 1:1 with isopropanol then applied to a Qiagen RNeasy Mini Kit (74106) and RNA purification was performed as per manufacturer’s directions. Library preparation was performed using a Kapa Biosystems mRNA Hyper Prep Kit. Sequencing was performed at the Vincent J Coates Genomic Sequencing Core at the University of California, Berkeley using an Illumina HS4000 mode SR100. Three biological replicates were measured per condition. Reads per gene were quantified using kallisto [43], with WBcel235 as the worm reference genome. Fold changes were determined using DESeq2 [44].

### Lifespan measurements

For lifespan experiments, animals were grown on either 2% or 4% RNAi plates on either EV or RNAi bacteria from L1 (see bleaching). Animals were then moved to their respective plates containing 100 µL of 100 mg/mL FUDR starting from day 1 of adulthood to eliminate progeny. For stress assays, animals were moved onto plates supplemented with either 2.5 mM paraquat or 25 µg/mL tunicamycin directly in the plate. Animals were grown at 20 °C and checked every 2 days for viability. For thermotolerance, day 1 adult animals were placed at 34 °C and were scored for viability every 2 hours. Animals were scored as dead if they did not exhibit any movement when prodded with a platinum wire at both the head and the tail. Animals that exhibited intestinal leakage, bagging, desiccation on the side of the petri dish, or other deaths unrelated to aging were scored as censored. All lifespan assay survival curves were plotted using Prism 7 Software and statistics were performed using Log-Rank testing. All lifespans were performed on 3 biological replicates and a table of lifespans is available in **Table S1**.

### C. elegans motility

For all motility experiments, thrashing of animals were scored on day 1, day 5, and day 9 animals grown on either 2% or 4% RNAi plates from L1. Animals were collected using M9 and placed onto an NGM plate without bacteria and 10 s video recordings were taken using a Leica M205FCA stereomicroscope. Thrashing was manually recorded wherein a single body bend in one direction is scored as a single thrash and the total number of thrashes for a 10 s period are calculated. Thrashing assays were performed on three biological replicates and statistics were calculated using Mann-Whitney testing using Prism 7 software.

### *C. elegans* reproduction assay

For all reproduction experiments, animals were grown on either 2% or 4% RNAi plates from L1 (see bleaching). Animals were singled out on the L4 stage and moved daily onto a new plate. Number of viable progeny on each plate was scored every day and cumulated per animal to calculate the number of viable progeny produced by each animal. Three biological replicates were performed on n > 10 animals were replicate and statistics were calculated using Mann-Whitney testing.

### Measurement of surface stiffness

Viscoelastic properties of NGM plates using agar from different commercially available sources were determined using an oscillatory rheometer (Anton-Paar, M302e) with parallel-plate geometry (8 mm) and a gap height of ∼ 0.2 mm under 0.1% strain and 1 Hz frequency at 37 °C in a humidity-controlled chamber.

## Author Contributions

R.H.S. designed and oversaw all experiments and methodology. M.O. and R.H.S. performed all experiments and prepared the manuscripts and figures. A.H. assisted in experimental design, RNA-seq analysis, and manuscript and figure preparation. M.A. and N.D. performed all high resolution microscopy. A.A. prepared all medium preparation and assisted in experimentation. T.C.T. and D.M. assisted in measurements of healthspan. M.V., A.B., A.J.C., D.H., A.W., T.W., and S.H. assisted in stress assays, K.M.T. performed all AFM experiments and data analysis, and G.G. performed RNA-seq analysis and assisted in experimental design and project management.

## DATA AVAILABILITY

All data required to evaluate the conclusions in this manuscript are available within the manuscript and Supporting Information. All strains synthesized in this manuscript are derivatives of N2 or other strains from CGC and are either made available on CGC or available upon request. All raw RNA-seq datasets are available through Annotare 2.0 Array Express Accession E-MTAB-13938.

## Acknowledgements

A.H., M.A., and G.G. are supported by T32AG052374; M.O., T.C.T., and M.V. are supported by 1R25AG076400 from the National Institute on Aging; T.W. is supported by the CIRM COMPASS Award EDUC5-13853; and R.H.S. is supported by R00AG065200 from the National Institute on Aging, Larry L. Hillblom Foundation Grant 2022-A-010-SUP, and the Glenn Foundation for Medical Research and AFAR Grant for Junior Faculty Award. Some strains were provided by the CGC, which is funded by the NIH Office of Research Infrastructure Programs (P40 OD010440). Some gene analysis was performed using Wormbase, which is funded on a U41 grant HG002223.

## Supplemental Figures and Figure Legends

**Fig. S1.**
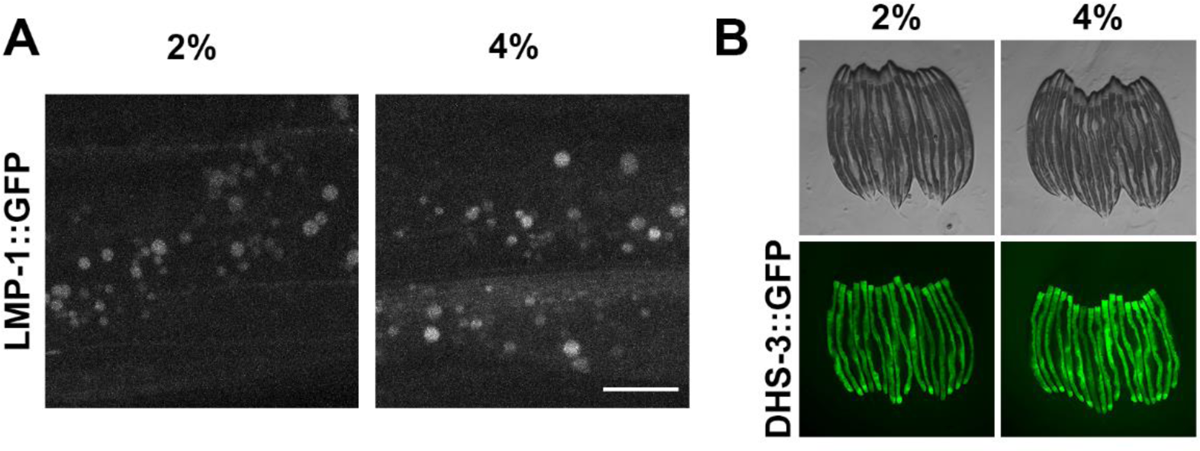
Growth on stiff substrates has no impact on lysosomes or lipid droplets. **(A)** Representative max projection fluorescent images of lysosomes by visualization of *lmp-1::GFP*. Images were captured on a Leica Stellaris system using optimized z-slices. **(B)** Representative fluorescent images of lipid droplets by visualization of DHS-3::GFP. For A-B, animals were grown on empty vector (EV) RNAi bacteria from L1 and imaged at day 1 of adulthood. Scale bar is 10 µm. Data is representative of 3 independent trials.

**Fig. S2.**
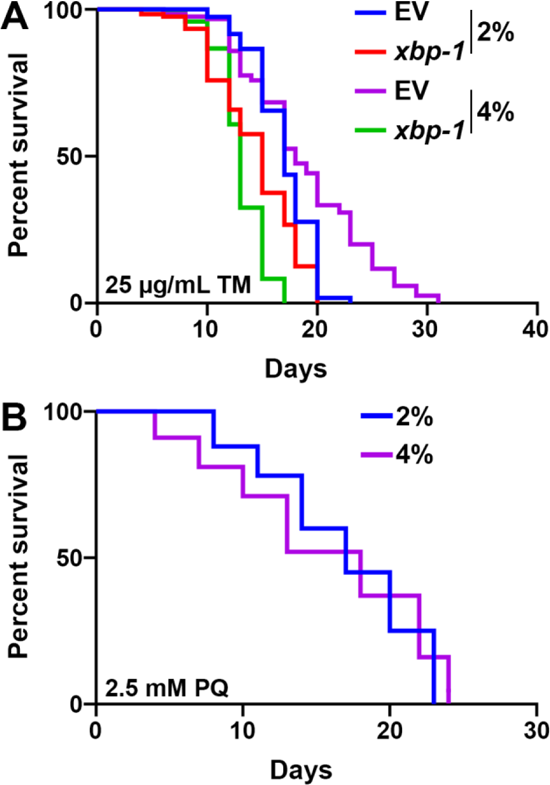
Growth on stiff substrates increases ER stress resilience. **(A)** N2 wild-type animals grown on empty vector (EV) or *xbp-1* RNAi on either control (2%) or stiff (4%) agar plates from L1. Animals were transferred onto the same RNAi and agar concentration plates containing 25 µg/mL tunicamycin (TM). Lifespans were scored every 2 days. Data is representative of 3 replicates and statistical analysis is available in **Table S1**. **(B)** N2 wild-type animals grown on EV RNAi on either control (2%) or stiff (4%) agar plates from L1. Animals were moved to plates containing 2.5 mM paraquat (PQ) at day 1 of adulthood and survival was scored every 2 days. Data is representative of 3 replicates and statistical analysis is available in **Table S1**.

**Fig. S3.**
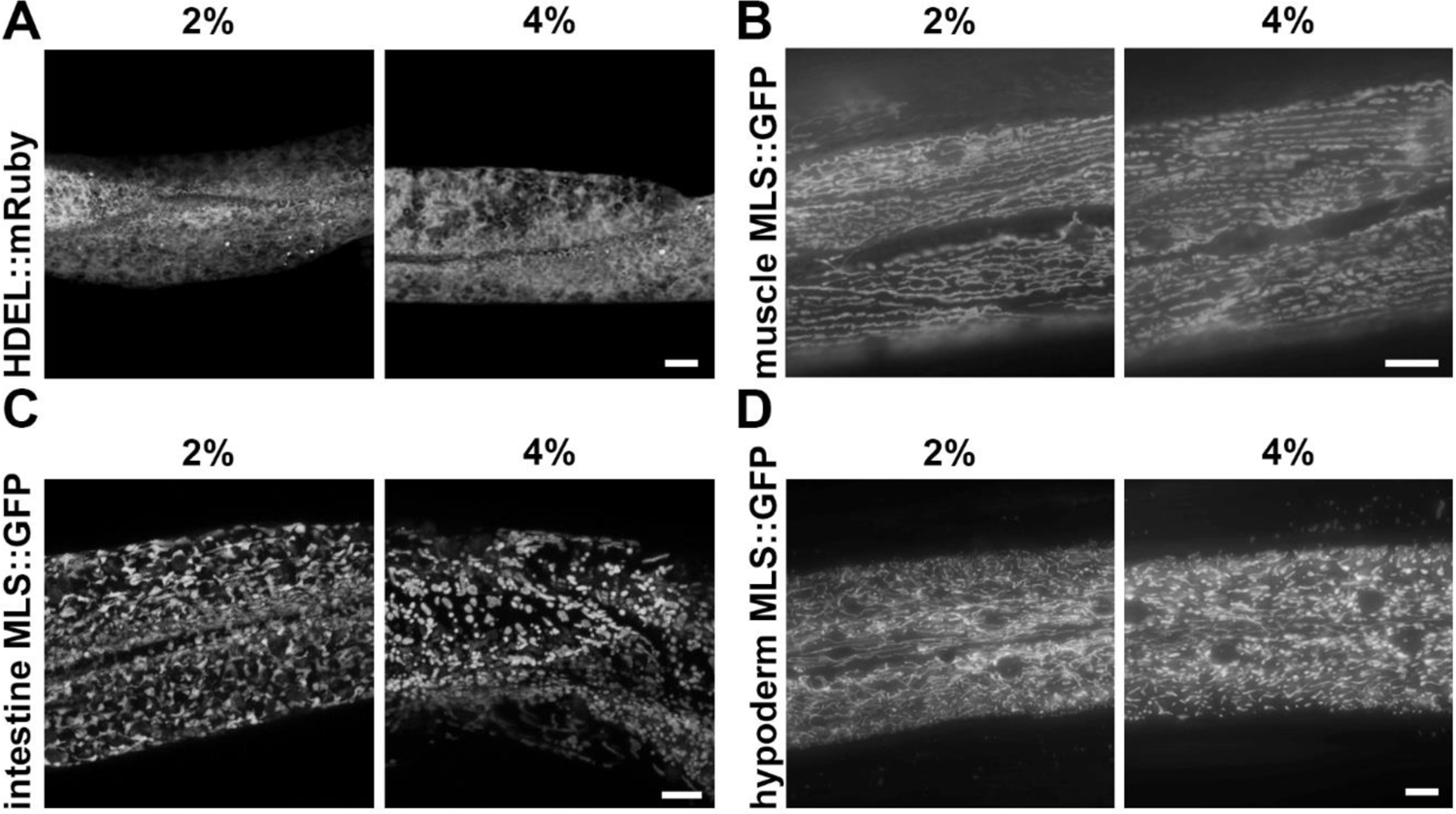
Growth on stiff substrates does not impact ER or mitochondrial structure. **(A)** Representative max projection fluorescent images of the endoplasmic reticulum in the intestine by visualization of *vha-6p::HDEL::mRuby*. Images were captured on a Leica Stellaris system using optimized z-slices. Animals were grown on empty vector (EV) bacteria from L1. Animals were moved onto 25 µg/mL tunicamycin containing plates at L4 and imaged at day 1 of adulthood. **(B)** Representative fluorescent images of body wall muscle mitochondria (*myo-3p::MLS::GFP*) are shown. Animals were grown on empty vector (EV) RNAi from L1. Single-slice images were captured on a Leica THUNDER Imager. Scale bar is 10 µm. **(C)** Representative max-projection fluorescent images of intestinal mitochondria (*gly-19p::MLS::GFP*) are shown. Animals were grown on empty vector (EV) RNAi from L1. Z-stack images were captured on a Leica THUNDER Imager using system-optimized z-slices. Scale bar is 10 µm. **(D)** Representative max-projection fluorescent images of hypodermal mitochondria (*col-19p::MLS::GFP*) are shown. Animals were grown on empty vector (EV) RNAi from L1. Z-stack images were captured on a Leica Stellaris using system-optimized z-slices. Scale bar is 10 µm.

**Fig. S4.**
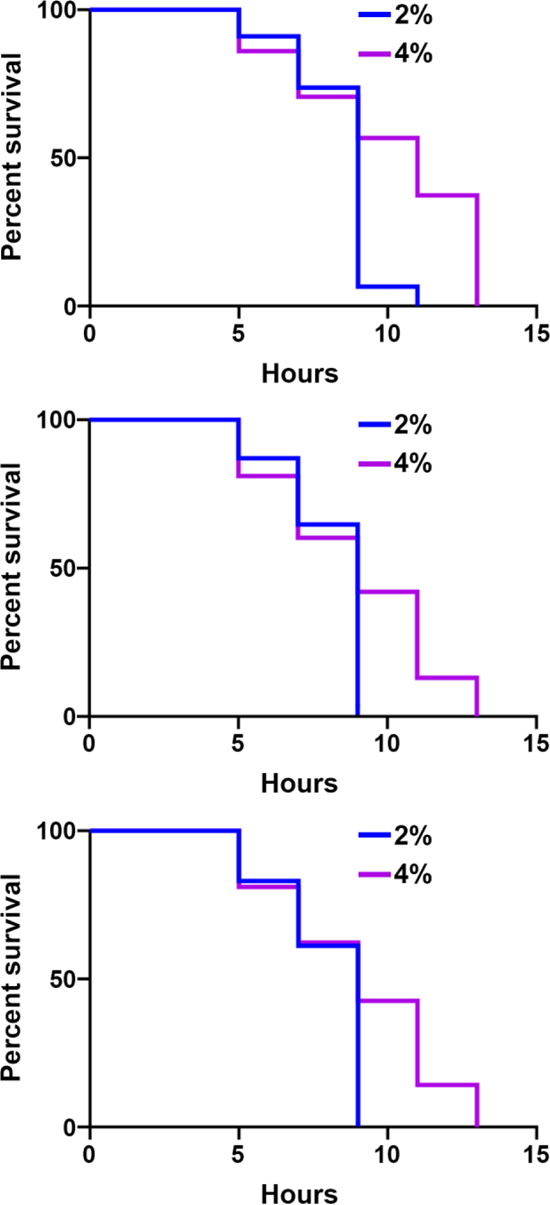
Growth on stiff substrates results in a mild increase in thermotolerance. N2 wild-type animals grown on EV RNAi on either control (2%) or stiff (4%) agar plates from L1. Animals were moved to 34°C at day 1 of adulthood and survival was scored every 2 hours. All three biological replicates are shown to highlight differences in maximal thermotolerance, despite similarities in median thermotolerance in some replicates. All statistics are available in **Table S1**.

## References

1. Levy Nogueira M, da Veiga Moreira J, Baronzio GF, Dubois B, Steyaert J-M, Schwartz L. Mechanical Stress as the Common Denominator between Chronic Inflammation, Cancer, and Alzheimer’s Disease. Front Oncol. 2015;5: 197. doi:10.3389/fonc.2015.00197

2. Galloway MT, Lalley AL, Shearn JT. The Role of Mechanical Loading in Tendon Development, Maintenance, Injury, and Repair. J Bone Joint Surg Am. 2013;95: 1620– 1628. doi:10.2106/JBJS.L.01004

3. Yi B, Xu Q, Liu W. An overview of substrate stiffness guided cellular response and its applications in tissue regeneration. Bioact Mater. 2021;15: 82–102. doi:10.1016/j.bioactmat.2021.12.005

4. Wang N. Review of Cellular Mechanotransduction. J Phys D Appl Phys. 2017;50: 233002.

5. Martino F, Perestrelo AR, Vinarský V, Forte G. Cellular Mechanotransduction: From Tension to Function. Front Physiol. 2018;9. doi:10.3389/fphys.2018.00824

6. Selman M, Pardo A. Fibroageing: An ageing pathological feature driven by dysregulated extracellular matrix-cell mechanobiology. Ageing Research Reviews. 2021;70: 101393. doi:10.1016/j.arr.2021.101393

7. Kwan KYC, Ng KWK, Rao Y, Zhu C, Qi S, Tuan RS, et al. Effect of Aging on Tendon Biology, Biomechanics and Implications for Treatment Approaches. Int J Mol Sci. 2023;24: 15183. doi:10.3390/ijms242015183

8. Lazarczuk SL, Maniar N, Opar DA, Duhig SJ, Shield A, Barrett RS, et al. Mechanical, Material and Morphological Adaptations of Healthy Lower Limb Tendons to Mechanical Loading: A Systematic Review and Meta-Analysis. Sports Med. 2022;52: 2405–2429. doi:10.1007/s40279-022-01695-y

9. Amargant F, Manuel SL, Tu Q, Parkes WS, Rivas F, Zhou LT, et al. Ovarian stiffness increases with age in the mammalian ovary and depends on collagen and hyaluronan matrices. Aging Cell. 2020;19: e13259. doi:10.1111/acel.13259

10. Papakonstantinou E, Roth M, Karakiulakis G. Hyaluronic acid: A key molecule in skin aging. Dermatoendocrinol. 2012;4: 253–258. doi:10.4161/derm.21923

11. Schinzel RT, Higuchi-Sanabria R, Shalem O, Moehle EA, Webster BM, Joe L, et al. The Hyaluronidase, TMEM2, Promotes ER Homeostasis and Longevity Independent of the UPRER. Cell. 2019;179: 1306–1318.e18. doi:10.1016/j.cell.2019.10.018

12. Wullkopf L, West A-KV, Leijnse N, Cox TR, Madsen CD, Oddershede LB, et al. Cancer cells’ ability to mechanically adjust to extracellular matrix stiffness correlates with their invasive potential. Mol Biol Cell. 2018;29: 2378–2385. doi:10.1091/mbc.E18-05-0319

13. Tharp KM, Higuchi-Sanabria R, Timblin GA, Ford B, Garzon-Coral C, Schneider C, et al. Adhesion-mediated mechanosignaling forces mitohormesis. Cell Metab. 2021; S1550–4131(21)00183–2. doi:10.1016/j.cmet.2021.04.017

14. Parida L, Ghosh UU, Padmanabhan V. The effects of groove height and substrate stiffness on *C. elegans* locomotion. Journal of Biomechanics. 2017;55: 34–40. doi:10.1016/j.jbiomech.2017.02.015

15. Castro Torres T, Moaddeli D, Averbukh M, Coakley AJ, Dutta N, Garcia G, et al. Surveying Low-Cost Methods to Measure Lifespan and Healthspan in Caenorhabditis elegans. J Vis Exp. 2022. doi:10.3791/64091

16. Durieux J, Wolff S, Dillin A. The cell-non-autonomous nature of electron transport chain-mediated longevity. Cell. 2011;144: 79–91. doi:10.1016/j.cell.2010.12.016

17. Taylor RC, Dillin A. XBP-1 is a cell-nonautonomous regulator of stress resistance and longevity. Cell. 2013;153: 1435–1447. doi:10.1016/j.cell.2013.05.042

18. Morley JF, Morimoto RI. Regulation of longevity in Caenorhabditis elegans by heat shock factor and molecular chaperones. Mol Biol Cell. 2004;15: 657–664. doi:10.1091/mbc.E03-07-0532

19. Özbey NP, Imanikia S, Krueger C, Hardege I, Morud J, Sheng M, et al. Tyramine Acts Downstream of Neuronal XBP-1s to Coordinate Inter-tissue UPRER Activation and Behavior in C. elegans. Developmental Cell. 2020;55: 754–770.e6. doi:10.1016/j.devcel.2020.10.024

20. Yoneda T, Benedetti C, Urano F, Clark SG, Harding HP, Ron D. Compartment-specific perturbation of protein handling activates genes encoding mitochondrial chaperones. J Cell Sci. 2004;117: 4055–4066. doi:10.1242/jcs.01275

21. Calfon M, Zeng H, Urano F, Till JH, Hubbard SR, Harding HP, et al. IRE1 couples endoplasmic reticulum load to secretory capacity by processing the XBP-1 mRNA. Nature. 2002;415: 92–96. doi:10.1038/415092a

22. Link CD, Cypser JR, Johnson CJ, Johnson TE. Direct observation of stress response in Caenorhabditis elegans using a reporter transgene. Cell Stress Chaperones. 1999;4: 235–242. doi:10.1379/1466-1268(1999)004<0235:doosri>2.3.co;2

23. Link CD, Johnson CJ. Reporter transgenes for study of oxidant stress in Caenorhabditis elegans. Methods Enzymol. 2002;353: 497–505. doi:10.1016/s0076-6879(02)53072-x

24. O’Rourke EJ, Ruvkun G. MXL-3 and HLH-30 transcriptionally link lipolysis and autophagy to nutrient availability. Nat Cell Biol. 2013;15: 668–676. doi:10.1038/ncb2741

25. Staab TA, McIntyre G, Wang L, Radeny J, Bettcher L, Guillen M, et al. The lipidomes of C. elegans with mutations in asm-3/acid sphingomyelinase and hyl-2/ceramide synthase show distinct lipid profiles during aging. Aging (Albany NY). 2023;15: 650–674. doi:10.18632/aging.204515

26. Treusch S, Knuth S, Slaugenhaupt SA, Goldin E, Grant BD, Fares H. Caenorhabditis elegans functional orthologue of human protein h-mucolipin-1 is required for lysosome biogenesis. Proc Natl Acad Sci U S A. 2004;101: 4483–4488. doi:10.1073/pnas.0400709101

27. Zhang P, Na H, Liu Z, Zhang S, Xue P, Chen Y, et al. Proteomic study and marker protein identification of Caenorhabditis elegans lipid droplets. Mol Cell Proteomics. 2012;11: 317–328. doi:10.1074/mcp.M111.016345

28. Metcalf MG, Higuchi-Sanabria R, Garcia G, Tsui CK, Dillin A. Beyond the cell factory: Homeostatic regulation of and by the UPRER. Science Advances. 2020;6: eabb9614. doi:10.1126/sciadv.abb9614

29. Castello PR, Drechsel DA, Patel M. Mitochondria are a major source of paraquat-induced reactive oxygen species production in the brain. J Biol Chem. 2007;282: 14186–14193. doi:10.1074/jbc.M700827200

30. Daniele JR, Higuchi-Sanabria R, Durieux J, Monshietehadi S, Ramachandran V, Tronnes SU, et al. UPRER promotes lipophagy independent of chaperones to extend life span. Science Advances. 2020;6: eaaz1441. doi:10.1126/sciadv.aaz1441

31. Moehle EA, Higuchi-Sanabria R, Tsui CK, Homentcovschi S, Tharp KM, Zhang H, et al. Cross-species screening platforms identify EPS-8 as a critical link for mitochondrial stress and actin stabilization. Sci Adv. 2021;7: eabj6818. doi:10.1126/sciadv.abj6818

32. Daniele JR, Esping DJ, Garcia G, Parsons LS, Arriaga EA, Dillin A. “High-Throughput Characterization of Region-Specific Mitochondrial Function and Morphology.” Sci Rep. 2017;7: 6749. doi:10.1038/s41598-017-05152-z

33. Higuchi-Sanabria R, Paul Rd JW, Durieux J, Benitez C, Frankino PA, Tronnes SU, et al. Spatial regulation of the actin cytoskeleton by HSF-1 during aging. Mol Biol Cell. 2018;29: 2522–2527. doi:10.1091/mbc.E18-06-0362

34. Egge N, Arneaud SLB, Wales P, Mihelakis M, McClendon J, Fonseca RS, et al. Age-Onset Phosphorylation of a Minor Actin Variant Promotes Intestinal Barrier Dysfunction. Developmental Cell. 2019;51: 587–601.e7. doi:10.1016/j.devcel.2019.11.001

35. Garcia G, Bar-Ziv R, Averbukh M, Dasgupta N, Dutta N, Zhang H, et al. Large-scale genetic screens identify BET-1 as a cytoskeleton regulator promoting actin function and life span. Aging Cell. 2023;22: e13742. doi:10.1111/acel.13742

36. Baird NA, Douglas PM, Simic MS, Grant AR, Moresco JJ, Wolff SC, et al. HSF-1-mediated cytoskeletal integrity determines thermotolerance and life span. Science. 2014;346: 360–363. doi:10.1126/science.1253168

37. Hoon JL, Tan MH, Koh C-G. The Regulation of Cellular Responses to Mechanical Cues by Rho GTPases. Cells. 2016;5: 17. doi:10.3390/cells5020017

38. Bar-Ziv R, Dutta N, Hruby A, Sukarto E, Henderson HR, Durieux J, et al. Glial-derived mitochondrial signals impact neuronal proteostasis and aging. bioRxiv; 2023. p. 2023.07.20.549924. doi:10.1101/2023.07.20.549924

39. Imanikia S, Özbey NP, Krueger C, Casanueva MO, Taylor RC. Neuronal XBP-1 Activates Intestinal Lysosomes to Improve Proteostasis in C. elegans. Curr Biol. 2019;29: 2322–2338.e7. doi:10.1016/j.cub.2019.06.031

40. Qu M, Zhang X, Hu X, Dong M, Pan X, Bian J, et al. BRD4 inhibitor JQ1 inhibits and reverses mechanical injury-induced corneal scarring. Cell Death Discov. 2018;4: 1–11. doi:10.1038/s41420-018-0066-1

41. Zhubanchaliyev A, Temirbekuly A, Kongrtay K, Wanshura LC, Kunz J. Targeting Mechanotransduction at the Transcriptional Level: YAP and BRD4 Are Novel Therapeutic Targets for the Reversal of Liver Fibrosis. Front Pharmacol. 2016;7: 462. doi:10.3389/fphar.2016.00462

42. Higuchi-Sanabria R, Durieux J, Kelet N, Homentcovschi S, de Los Rios Rogers M, Monshietehadi S, et al. Divergent Nodes of Non-autonomous UPRER Signaling through Serotonergic and Dopaminergic Neurons. Cell Rep. 2020;33: 108489. doi:10.1016/j.celrep.2020.108489

43. Bray NL, Pimentel H, Melsted P, Pachter L. Near-optimal probabilistic RNA-seq quantification. Nat Biotechnol. 2016;34: 525–527. doi:10.1038/nbt.3519

44. Love MI, Huber W, Anders S. Moderated estimation of fold change and dispersion for RNA-seq data with DESeq2. Genome Biology. 2014;15: 550. doi:10.1186/s13059-014-0550-8

